# Gallocin A, an atypical two-peptide bacteriocin with intramolecular disulfide bonds required for activity

**DOI:** 10.1101/2022.12.05.519244

**Authors:** Alexis Proutière, Laurence du Merle, Marta Garcia-Lopez, Corentin Léger, Alexis Voegele, Alexandre Chenal, Antony Harrington, Yftah Tal-Gan, Thomas Cokelaer, Patrick Trieu-Cuot, Shaynoor Dramsi

**Author notes:** Present address: Laboratory of Molecular Microbiology, Global Health Institute, School of Life Sciences, Ecole Polytechnique Fédérale de Lausanne (EPFL), Lausanne, Switzerland. Correspondence: Shaynoor Dramsi, or Alexis Proutière.

## Abstract

*Streptococcus gallolyticus* subsp. *gallolyticus* (*SGG*) is an opportunistic gut pathogen associated with colorectal cancer. We previously showed that colonization of the murine colon by *SGG* in tumoral conditions was strongly enhanced by the production of gallocin A, a two-peptide bacteriocin. Here, we aimed at characterizing the mechanisms of its action and resistance. Using a genetic approach, we demonstrated that gallocin A is composed of two peptides, GllA1 and GllA2, which are inactive alone and act together to kill “target” bacteria. We showed that gallocin A can kill phylogenetically close relatives. Importantly, we demonstrated that gallocin A peptides can insert into membranes and permeabilize lipid bilayer vesicles. Next, we showed that the third gene of the gallocin A operon named GIP, is necessary and sufficient to confer immunity to gallocin A. Structural modelling of GllA1 and GllA2 mature peptides suggested that both peptides form alpha-helical hairpins stabilized by intramolecular disulfide bridges. The presence of a disulfide bond in GllA1 and GllA2 was confirmed experimentally. Addition of disulfide reducing agents abrogated gallocin A activity. Likewise, deletion of a gene encoding a surface protein with a thioredoxin-like domain impaired gallocin A ability to kill *Enterococcus faecalis*. Structural modelling of GIP revealed a hairpin-like structure strongly resembling that of the GllA1 and GllA2 mature peptides, suggesting a mechanism of immunity by competition with GllA1/2. Finally, identification of other class IIb bacteriocins exhibiting a similar alpha-helical hairpin fold stabilized with an intramolecular disulfide bridge suggests the existence of a new subclass of class IIb bacteriocins.

**IMPORTANCE:** *Streptococcus gallolyticus subsp. gallolyticus* (*SGG*), previously named *Streptococcus bovis* biotype I, is an opportunistic pathogen responsible for invasive infections (septicemia, endocarditis) in elderly people and often associated with asymptomatic colon tumors. *SGG* is one of the first bacteria to be associated with the occurrence of colorectal cancer in humans. Previously, we showed that tumor-associated conditions in the colon provide to *SGG* with the ideal environment to proliferate at the expense of phylogenetically and metabolically closely related commensal bacteria such as enterococci (Aymeric et al., 2017). *SGG* takes advantage of CRC-associated conditions to outcompete and substitute commensal members of the gut microbiota using a specific bacteriocin named gallocin and renamed gallocin A recently following the discovery of gallocin D in a peculiar *SGG* isolate. Here, we showed that gallocin A is a two-peptide bacteriocin and that both GllA1 and GllA2 peptides are required for antimicrobial activity. Gallocin A was shown to permeabilize bacterial membranes and to kill phylogenetically closely related bacteria such as most streptococci, lactococci and enterococci, probably through membrane pore formation. GllA1 and GllA2 secreted peptides are unusually long (42 and 60 amino acids long) and with very few charged amino acids compared to well-known class IIb bacteriocins. *In silico* modelling revealed that both GllA1 and GllA2 exhibit a similar hairpin-like conformation stabilized by an intramolecular disulfide bond. We also showed that the GIP immunity peptide also forms a hairpin like structure like GllA1/GllA2. Thus, we hypothesize that GIP blocks the formation of the GllA1/GllA2 complex by interacting with GllA1 or GllA2. Gallocin A may constitute the first class IIb bacteriocin displaying disulfide bridges important for its structure and activity and the founding member of a subtype of class IIb bacteriocins.

## INTRODUCTION

*Streptococcus gallolyticus subsp. gallolyticus* (*SGG*), formerly known as *S. bovis* biotype I, is a gut commensal of the rumen of herbivores causing infective endocarditis in elderly people and strongly associated with colorectal cancer (CRC). In a previous study, we have shown that *SGG* is able to take advantage of tumoral conditions (increased secondary bile salts concentration) to thrive and colonize the intestinal tract of Notch/APC mice. This colonization advantage was shown to be linked to the production of a two-component bacteriocin named gallocin enabling *SGG* to outcompete murine gut resident enterococci in tumor-bearing mice, but not in non-tumor mice (1). As such, gallocin constitutes the first bacterial factor explaining *SGG* association with CRC. Identification of a different gallocin, named gallocin D, from the environmental isolate *SGG* LL009 (2) led to renaming gallocin of *SGG* UCN34 as gallocin A.

Bacteriocins are highly diverse antimicrobial peptides secreted by nearly all bacteria. In gram-positive bacteria, they are divided in three classes based on size, amino acid composition and structure (3). Class I includes small (< 10-kDa), heat-stable peptides that undergo enzymatic modification during biosynthesis; class II includes small (< 10 kDa) heat-stable peptides without post-translational modifications; class III includes larger (> 10 kDa), thermo-labile peptides and proteins. Class II bacteriocins are further subdivided into four subtypes: class IIa consists of pediocin-like bacteriocins, class IIb consists of bacteriocins with two peptides, class IIc consists of leaderless bacteriocins, and class IId encompass all other non-pediocin-like, single peptide bacteriocins with a leader sequence. Previous *in silico* analysis revealed that gallocin A, encoded by *gallo*_*2021* (renamed *gllA2*) and *gallo_2020* (renamed *gllA1*), belong to the class IIb bacteriocins (Pfam10439) exhibiting a characteristic double glycine leader peptide. The third gene of this operon (*gallo_2019* renamed *gip*) was thought to encode the immunity protein.

We previously showed that a secreted peptide, GSP, activates transcription of the gallocin A core operon through a two-component system named BlpHR (4). The entire BlpHR regulon has been characterized and consists of 24 genes, of which 20 belong to the gallocin locus (4). Concomitantly, we showed that GSP but also GllA1 and GllA2 are secreted by a unique ABC transporter named BlpAB (5). GllA1 and GllA2 are synthesized as pre-peptides with an N-terminal leader sequence cleaved during export after a double glycine motif to produce the extracellular mature active peptide. Well-known class IIb bacteriocins are usually constituted of two genes encoding short peptides, named alpha and beta, that fold into alpha-helical structures and insert into target bacterial membranes to alter their permeability, resulting in ion leakage and cell death (6).

The aim of the present work was to characterize the gallocin A activity spectrum, its mode of action and the immunity mechanism. Our results indicate that GllA1 and GllA2 peptides are atypical and contain a disulfide bond required for antibacterial activity. We showed that GllA1/GllA2 can permeabilize lipid bilayers. The predicted structure of the GIP immunity peptide strikingly mimics that of the GllA1 and GllA2 mature peptides suggesting a mechanism of immunity by interference. *In vitro*, gallocin A was able to kill most closely related species such as streptococci and enterococci, highlighting the potential of these narrow-spectrum antimicrobials as alternatives to antibiotics.

## RESULTS

### Gallocin A is a two-peptide bacteriocin

As shown in Fig. 1A, the gallocin A core operon is composed of three genes (*gllA2, gllA1, gip*) coding for 2 putative bacteriocin peptides (GllA1 and GllA2) and a putative immunity protein (GIP). To demonstrate the role of *gllA1* and *gllA2* in gallocin A activity, we performed in-frame deletions of *gllA1* and *gllA2* separately in *SGG* strain UCN34 (wild-type, WT) and tested the antibacterial activity of the corresponding mutant supernatants by plate diffusion assays, as described previously (4). As shown in Fig. 1B, the antimicrobial activity of gallocin A is completely abolished in the supernatants of *ΔgllA1* and *ΔgllA2* mutants and restored when the supernatants of *ΔgllA1* and *ΔgllA2* are combined in a 1:1 ratio. This result demonstrates that both GllA1 and GllA2 are required for gallocin A activity and confirms that gallocin A is a two-peptide Class IIb bacteriocin (3). Finally, we showed that gallocin A is active in a wide range of pH (2-12, Fig. S1A) and temperature (Fig. S1B).

**Fig. 1.**
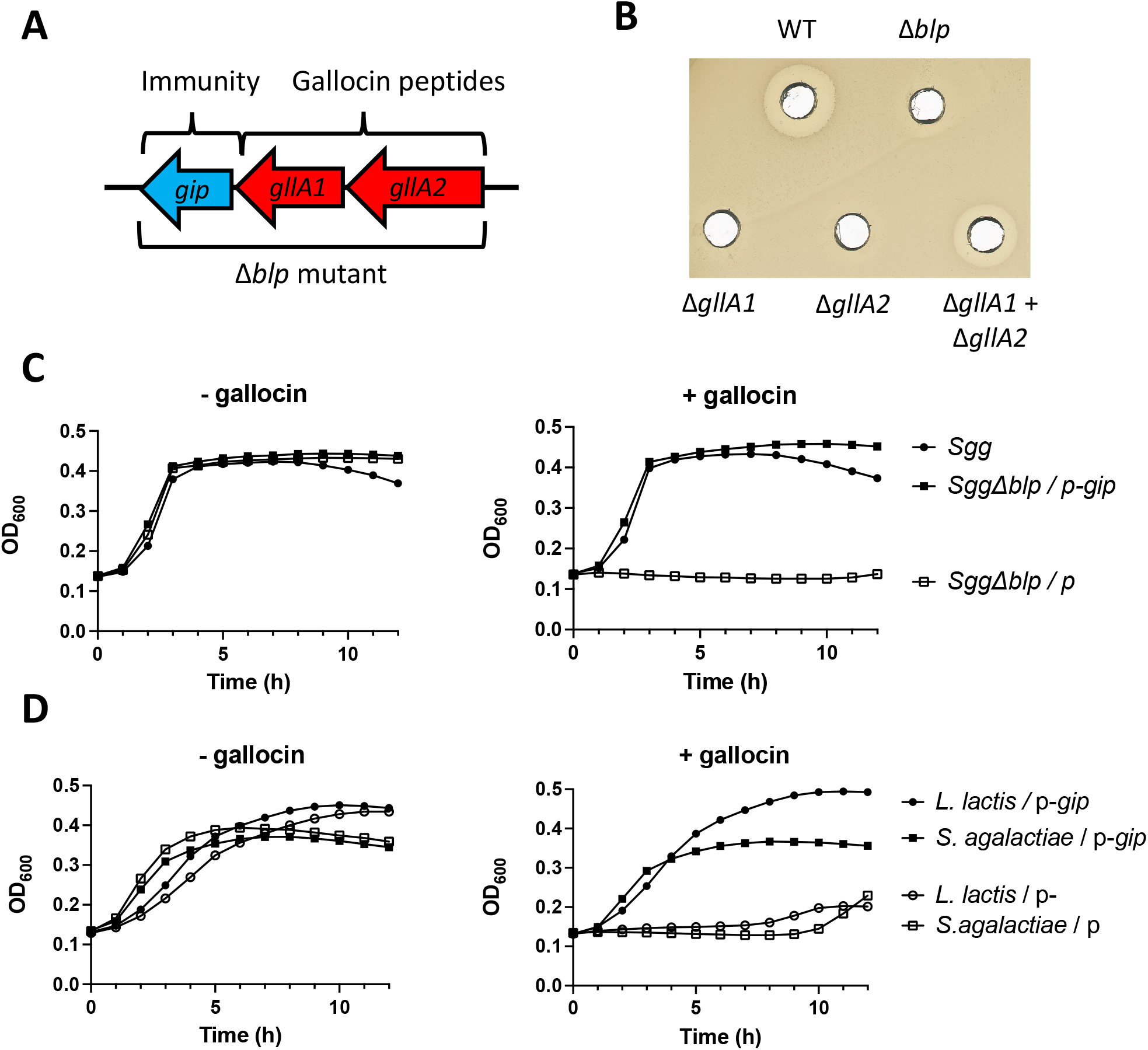
Gallocin A is a two-peptide bacteriocin. **A)** The core operon encoding gallocin A peptides and the immunity protein in SGG strain UCN34. Gallocin genes are indicated in red and renamed *gllA1* and *gllA2* according to (2). **B)** Agar diffusion assay to test gallocin activity from supernatants of UCN34 WT, Δ*gllA1*, Δ*gllA2* et Δ*blp* against gallocin-sensitive *S. gallolyticus subsp. macedonicus* (*SGM*) strain. **C)** and **D)** Growth curves of *SGG* Δ*blp, S. agalactiae* A909 and *L. lactis* NZ9000 containing an empty plasmid (p) or a plasmid expressing *gip* (p-*gip*) in THY supplemented with supernatant of Δ*blpS* (a strain overproducing gallocin, “+gallocin”) or Δ*blp* (gallocin deletion mutant, “-gallocin”) and 0.01% of tween 20.

Since the gene encoding the putative immunity protein named GIP cannot be deleted alone without self-intoxication of the bacteria, we used the original mutant UCN34*Δblp* (1) in which the three genes of gallocin A operon (*gllA2-gllA1-gip*) were deleted and tested its sensitivity to gallocin A. As expected, the Δ*blp* mutant became sensitive to gallocin A (Fig. 1C). Next, we complemented the *Δblp* mutant with a plasmid encoding *gip* and showed that this was sufficient to restore bacterial growth of the recombinant strain in the presence of gallocin A. These results demonstrate that GIP confers immunity to gallocin A (Fig. 1C). Moreover, constitutive expression of *gip* in heterologous bacteria sensitive to gallocin (such as *Streptococcus agalactiae* and *Lactococcus lactis*) allowed their growth in the presence of gallocin (Fig. 1D). These results clearly demonstrate that expression of *gip* alone is necessary and sufficient to confer full immunity against gallocin A.

### Gallocin A is active against various streptococci and enterococci

To further characterize the gallocin A activity spectrum, we tested the sensitivity of various bacteria from our laboratory collection, including species found as commensals in the gut as well as known gram-positive human pathogens. We showed that gallocin A is active only against closely related bacteria, including various streptococci, enterococci, lactococci and inactive against all other gram-positive and gram-negative bacteria tested (Fig. 2, Fig. S2A). Interestingly, the three different *S. agalactiae* strains tested (NEM316, BM110, and A909) differed significantly in their susceptibility to gallocin A. Similarly, sensitivity to gallocin A of many *Enterococcus faecalis* clinical isolates, including a few vancomycin resistant isolates, was also variable (Fig. S2B). These results indicate that gallocin A sensitivity of a given species can vary from one strain to another.

**Fig. 2.**
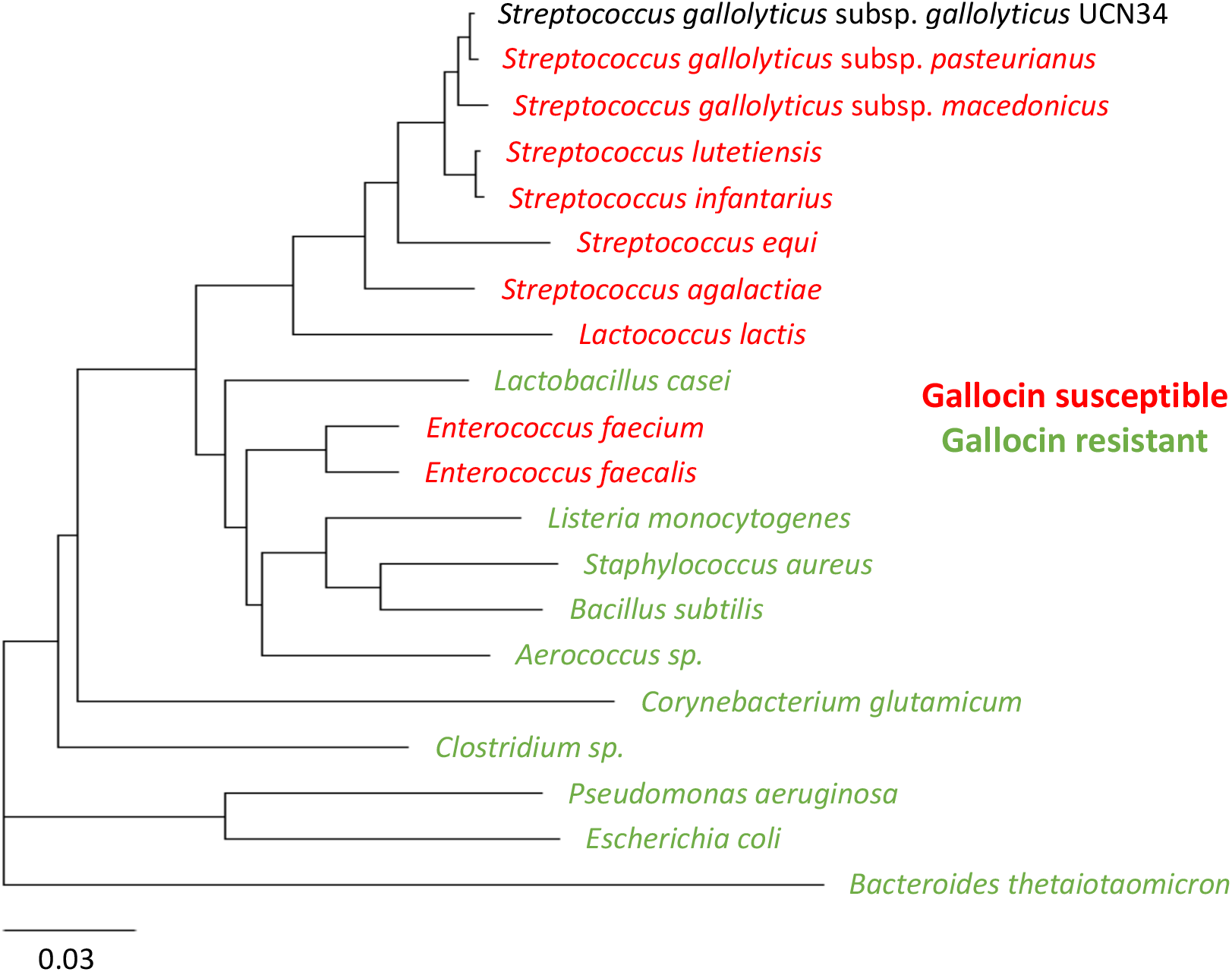
Gallocin A is active against most streptococci, lactococci and enterococci. Phylogenetic tree based on the 16S RNA sequence (from the Silva online database) of different bacterial species that are resistant (in red) or sensitive (in green) to gallocin, as determined by agar diffusion assay (Fig. S2).

### Gallocin A induces target cell-membrane depolarization

To test whether gallocin A peptides can alter cell membrane permeability, as shown for well-studied class IIb bacteriocins, we assessed its impact on target cell membrane potential using the fluorescent voltage-dependent dye DiBAC4(3) and propidium iodide (PI). DiBAC4(3) can access the cytoplasm only when the membrane is depolarized, thus indicating an ion imbalance, and the DNA intercalator PI can only enter bacterial cells when the cytoplasmic membrane is compromised. The entry of PI and DiBAC4(3) in cells exposed to supernatants from UCN34 WT, Δ*blp* (no gallocin A) and Δ*blpS* (a mutant previously shown to overproduce gallocin A, (4)) was assessed by flow cytometry. As shown in Fig. 3A and B, fluorescent dye penetration in *E. faecalis* OG1RF was increased in the presence of gallocin A as compared to the control supernatant without gallocin A, indicating that gallocin A peptides can form pores in bacterial membranes.

**Fig. 3.**
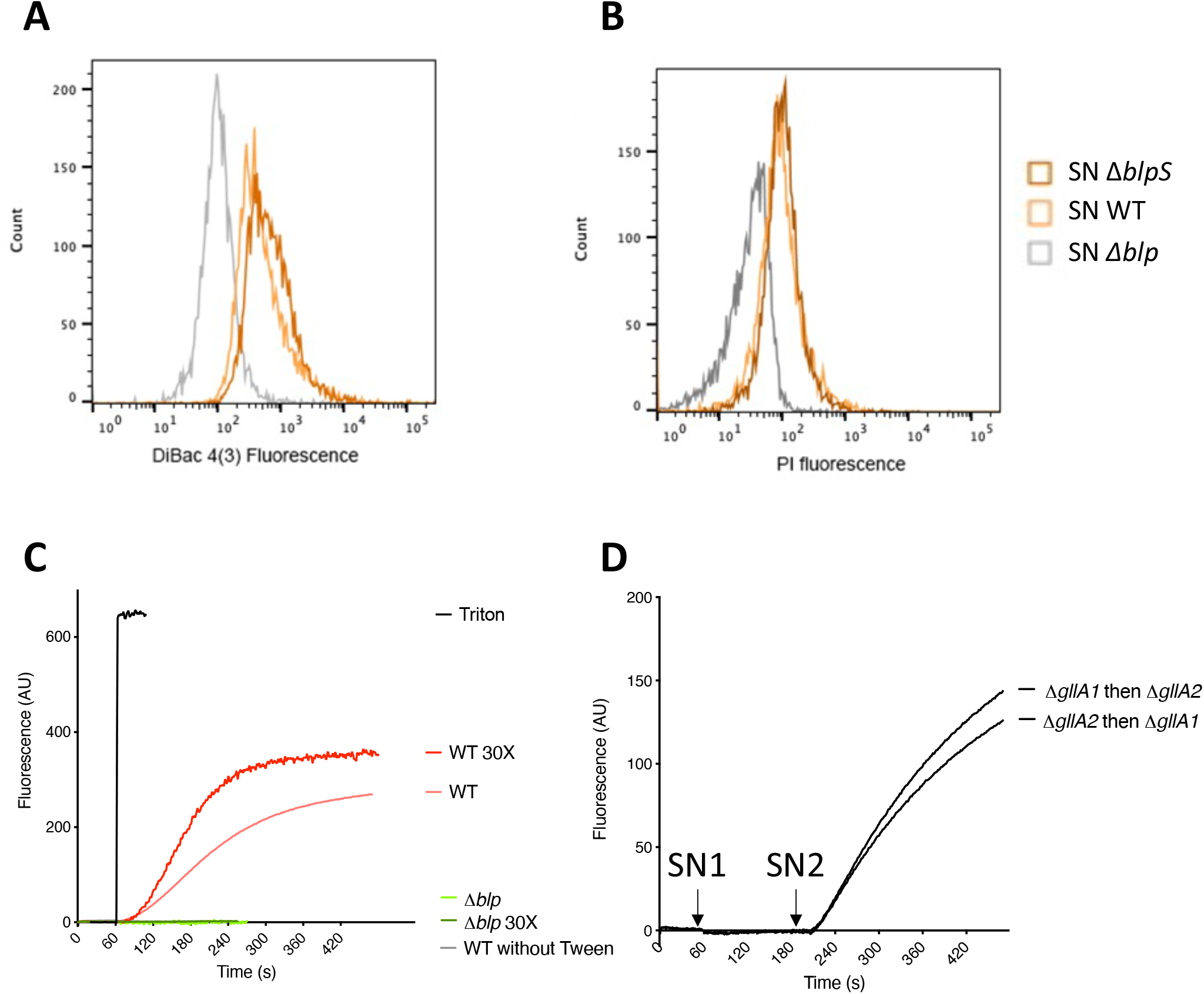
Gallocin A can permeabilize bacterial membranes and lipid vesicles. Fluorescence of the voltage-sensitive DiBac4(3) **(A)** or the membrane impermeant propidium iodide PI **(B)** after resuspension of *Enterococcus faecalis* OG1RF in supernatant of UCN34 WT, Δ*blp* (-gallocin) and Δ*blpS* (overexpressing gallocin). **C-D)** Measure of the fluorescence corresponding to the release of ANTS (ex: 390nm, em: 515nm) encapsulated in large unilamellar vesicles after addition of *SGG* supernatant or Triton X-100 (positive control). **C:** At 60 s, Triton or the supernatant of *SGG* UCN34 WT, or Δ*blp*, or WT 30X (concentrated 30 times) or Δ*blp* 30X, were added to the liposomes. **D:** At 60 s (SN1), the supernatant of Δ*gllA1* or Δ*gllA2* was added to the lipid vesicle suspension. At 200 s (SN2), the supernatant of the other strain is added. AU: Arbitrary Unit.

It was previously shown that pore formation by the two-peptide bacteriocins lactococcin G and enterocin 1071 requires the presence of UppP, a membrane protein involved in peptidoglycan synthesis that could serve as a receptor for these bacteriocins (7). To investigate whether gallocin A is active in the absence of a proteinaceous receptor, we tested its capacity to permeabilize lipid bilayer vesicles. To do so, we used large unilamellar vesicles (LUV) in which a fluorescence marker, the 8-Aminonaphthalene-1,3,6-Trisulfonic Acid (ANTS) and its quencher, p-Xylene-Bis-Pyridinium Bromide (DPX), are encapsulated. If pores are formed in the membrane of the liposomes, ANTS and DPX are released in the medium and ANTS recovers its fluorescence. As shown in Fig. 3C, addition of UCN34 WT supernatant containing gallocin A led to LUV permeabilization while the supernatant of the Δ*blp* mutant had no effect, showing that gallocin A can alter the vesicle membrane. Of note, addition of small amount of Tween 20 (0.01%) was necessary to observe gallocin A activity. Importantly, the Δ*blp* supernatant supplemented Tween20 at 0.01% had no effect on liposomes, showing that the membrane permeabilization induced by the UCN34 WT supernatant is not caused by the detergent alone (Fig. 3C).

We also confirmed that both GllA1 and GllA2 were required for membrane permeabilization. Indeed, addition of Δ*gllA1* or Δ*gllA2* supernatant alone had no effect, while addition of both supernatants led to LUV permeabilization, regardless of which peptide was added first (Fig. 3D).

### Gallocin A peptides contain a disulfide bond essential for their bactericidal activity

Both GllA1 and GllA2 pre-peptides exhibit a typical N-terminal leader sequences of 23 amino acids, ending with two glycine residues, which is cleaved upon secretion of these peptides through a dedicated ABC transporter (5). GllA1 and GllA2 mature peptides each contain 2 cysteines, which can potentially form a disulfide bridge important for their structure and function. Indeed, we showed that addition of reducing agents such as dithiothreitol (DTT) or β-mercaptoethanol (data not shown) abolished gallocin A activity (Fig. 4A), whereas it has no effect on a control bacteriocin which does not possess a disulfide bond, such as nisin. Furthermore, LC/MS analysis provided the exact molecular masses of the mature GllA1 and GllA2 peptides. The calculated masses identified oxidized cysteine residues, indicating the presence of a disulfide bridge in each peptide (Fig. S3).

**Fig. 4.**
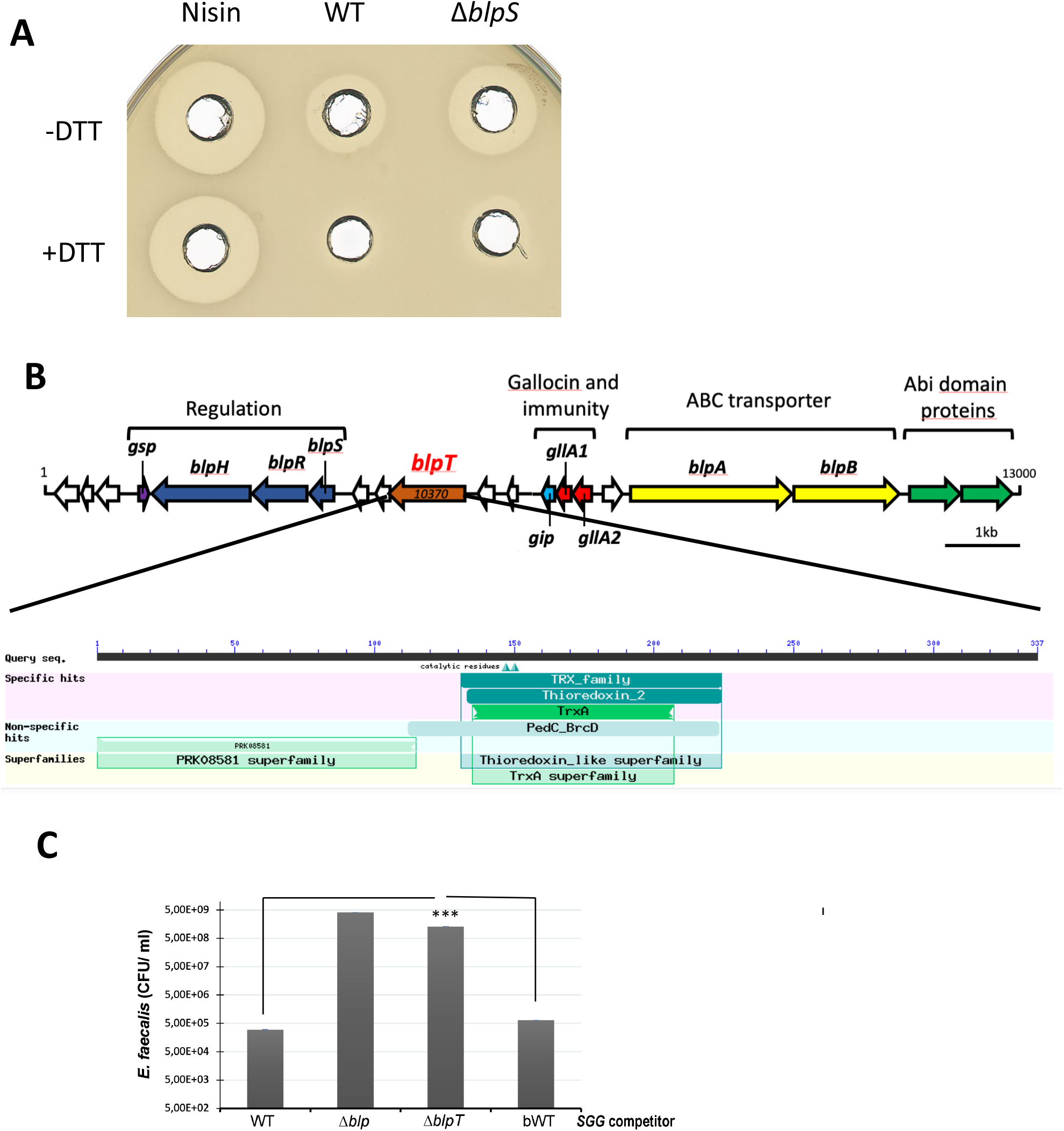
Gallocin A peptides possess a disulfide bridge important for structure and activity. **A)** Agar diffusion assay to test gallocin activity from supernatants of *SGG* WT or Δ*blpS* supplemented or not with 50mM of DTT **B)** Schematic representation of the gallocin genomic locus and pBLAST domain identification in BlpT protein. **C)** Recovered *E. faecalis* after co-culture at 1:1 ratio for 4 h with *SGG* WT, Δ*blp*, Δ*blpT* and WT revertant from *blpT* deletion.

Interestingly, the gallocin A genomic locus in *SGG* UCN34 contains a conserved co-regulated gene (4), *gallo_rs10370*, encoding a putative “bacteriocin biosynthesis protein” containing a thioredoxin domain (Fig. 4B). The thioredoxin domain is known to facilitate disulfide bond formation in *E. coli* (8) and is predicted to be extracellular by Pfam/Interproscan. We hypothesized that this gene renamed *blpT*, which encodes a surface protein potentially anchored to the cell-wall, could assist disulfide bond formation in gallocin A peptides following secretion and cleavage of the leader peptide by the ABC transporter BlpAB (5). Indeed, deletion of this gene in UCN34 (*ΔblpT*) strongly altered the ability of *SGG* to outcompete *Enterococcus faecalis* OG1RF in competition experiments where attacker *SGG* and prey *E. faecalis* were inoculated together in THY liquid medium at a 1:1 ratio and counted on entero-agar plates after 4 h of co-culture at 37°C (Fig. 4C). Remarkably, the Δ*blpT* mutant was comparable to the Δ*blp* mutant and the back to the WT behaved like the parental UCN34 WT (Fig. 4C). Altogether these results indicate the existence of disulfide bond in gallocin A mature peptides important for activity. Of note, the disulfide bond formation pathway of *E. coli*, containing the thioredoxin-like protein DsbA, was shown to be particularly important in anaerobic conditions (9). It is thus tempting to speculate that BlpT activity could be particularly important in the anaerobic environment that *SGG* encounter in the colon.

### The structural models of gallocin A peptides differ from those of other two-peptide bacteriocins

Structural modelling of GllA1 and GllA2 pre- and mature forms was performed using ColabFold (10) and showed that the putative N-terminal leader sequences adopt disordered and extended conformations (Fig. 5A and B). The structural models of mature GllA1 and GllA2 are composed of two antiparallel alpha-helices, i.e. adopting an alpha-helical hairpin fold (Fig. 5A and B and Fig. S4A and B). Interestingly, the two cysteines of GllA1 and GllA2 are facing one other in each alpha-helix of the helical hairpins, forming an intramolecular disulfide bond. This suggests that the disulfide bonds in GllA1 and GllA2 reduce the conformational flexibility within each alpha-helical hairpin and stabilize their three-dimensional structures. Interestingly, modelling of the immunity peptide GIP shows striking structural similarities with those of the mature GllA1 and GllA2 peptides (Fig. 6A and Fig. S4C). Despite a relative low confidence (lDDT between 50 and 65 %), the five structural models of GllA1/GllA2, GllA1/GIP and GllA2/GIP show similar orientations, giving credit to these models (Fig. 6B-D and Fig. S4D-F). As shown by aligning Cα of each GIP in the GllA1/GIP and GllA2/GIP, we hypothesized that GIP could intercalate between GllA1 and GllA2 (Fig. 6E). Thus, GIP might provide immunity by preventing interaction between GllA1 and GllA2 within the bacterial cell membrane of the producing bacteria.

**Fig. 5.**
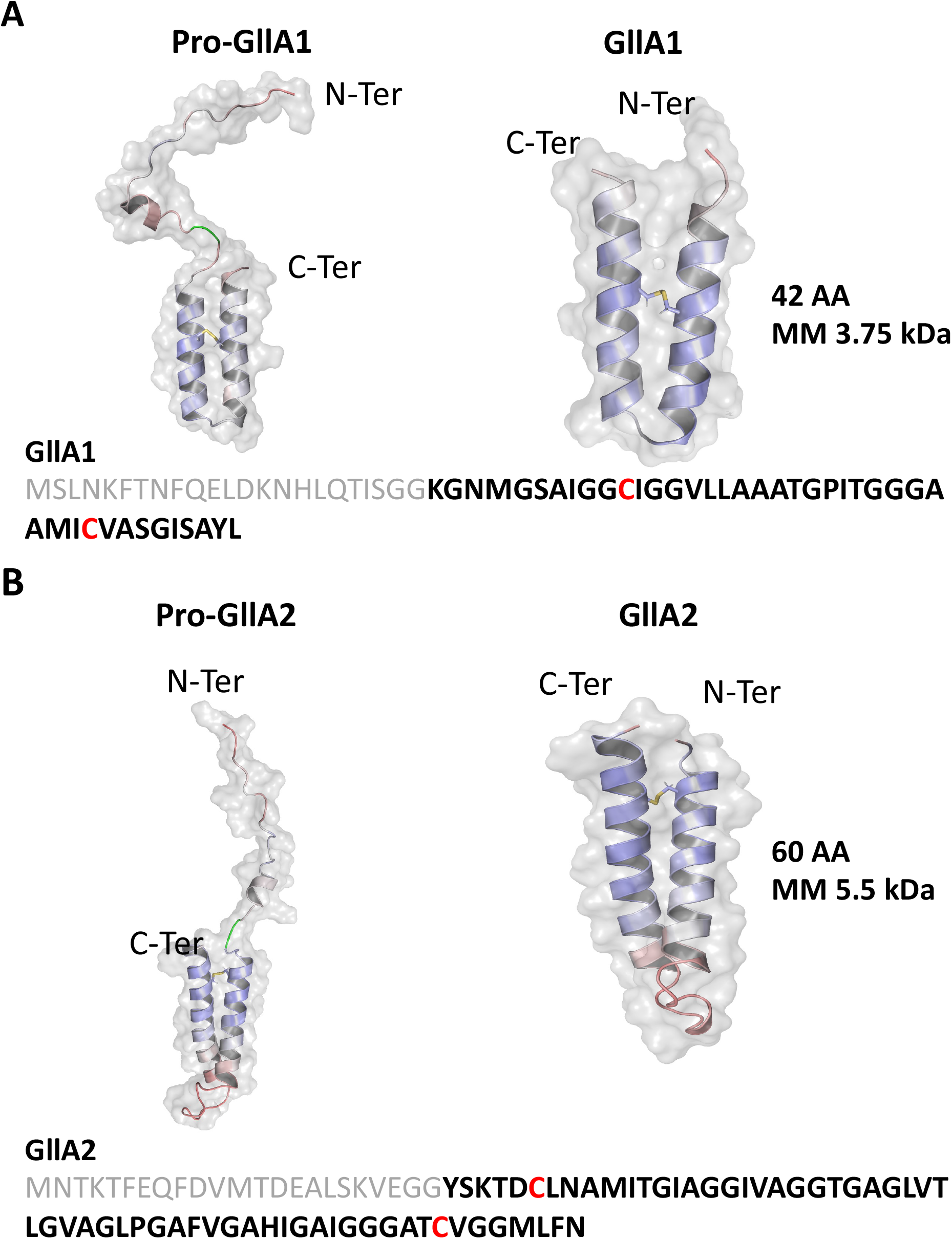
Structural models of GllA1 and GllA2 predicted using ColabFold. Pro- and mature forms of GllA1 (A) and GllA2 (B) using ColabFold and visualization was obtained with PyMOL (version 2.5.2 The PyMOL Molecular Graphics System, Version 2.0 Schrödinger, LLC). All representations are colored with predicted lDDT from a score of 30% (red) to 100% (blue). For the pro-GllA1 and pro-GllA2, glycine doublet is colored in green. The disulfide bridges are represented in stick.

**Fig. 6.**
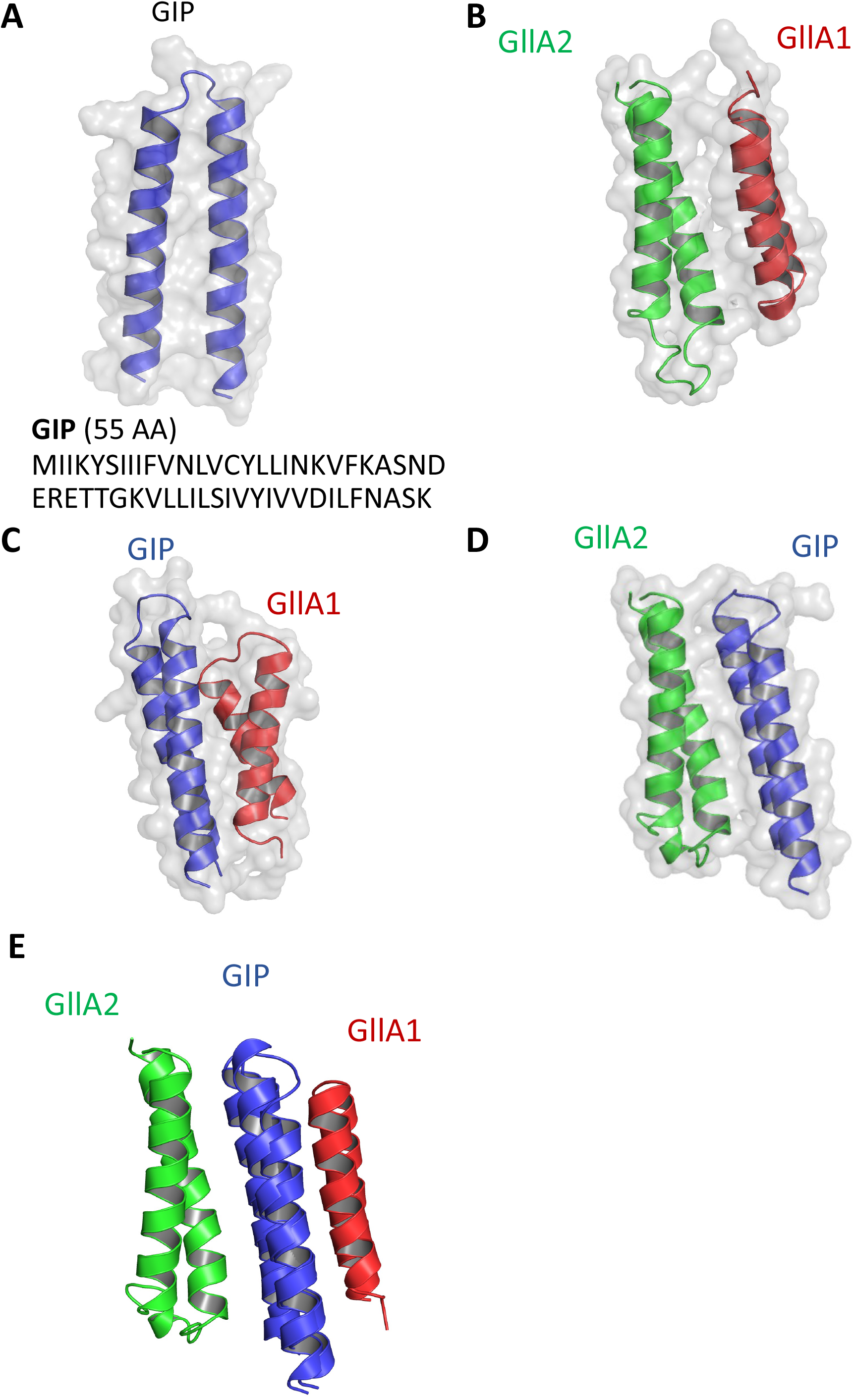
Structural models of GIP and its interactions with GllA1, GllA2. **A)** ColabFold modelling of GIP and visualization with PyMOL. **B, C, D)** ColabFold modelling of the interaction between GllA1/GllA2 **(B)**, GIP/GllA1 **(C)**, GIP/GllA2 **(D)**, and GllA1/GIP/GllA2 **(E)** interaction models aligned on the C⍰ of each GIP.

### Mechanisms of resistance to gallocin A

To better understand the mode of action of gallocin A, we decided to investigate the mechanisms of resistance to gallocin A. For that purpose, we isolated 14 spontaneous mutants (named RSM 1 to 14) of the highly sensitive strain *S. gallolyticus* subsp. *macedonicus* CIP 105683T on agar plates supplemented with gallocin A (see Materials and Methods for details). As shown in the supplemental Fig. S5B and C, 12 out of these 14 mutants were able to grow in liquid THY supplemented with gallocin A, in contrast to the parental strain *SGM* WT. However, when grown in presence of the control Δ*blp* supernatant, which does not contain gallocin A, all the mutants exhibited a longer latency phase than the parental *SGM* WT, suggesting that the acquired mutations may have a fitness cost.

To identify the mutations conferring resistance to gallocin A in these mutants, whole-genome sequencing was performed using Illumina technology and compared with the genome of the parental strain that was *de novo* assembled using PacBio sequencing. Between 1 and 8 single nucleotide polymorphism (SNP)/deletion/insertion were identified in each RSM mutant when compared to the WT controls (Table 1). Seven out of twelve mutants (RSM1, RSM2, RSM4, RSM5, RSM6, RSM12, RSM14) had mutations in the genes encoding the WalKR two-component system (TCS) and 3 others (RSM 7, RSM 8 and RSM10) had mutations in a gene (homologous to *gallo_rs1495*) encoding a putative “aggregation promoting factor” which contain a LysM peptidoglycan-binding domain and a lysozyme-like domain (Table 1, Fig. S6). The 2 remaining mutants (RSM3 and RSM11) displayed mutations which were not present in the other mutants and located in other genes.

**Table 1:**
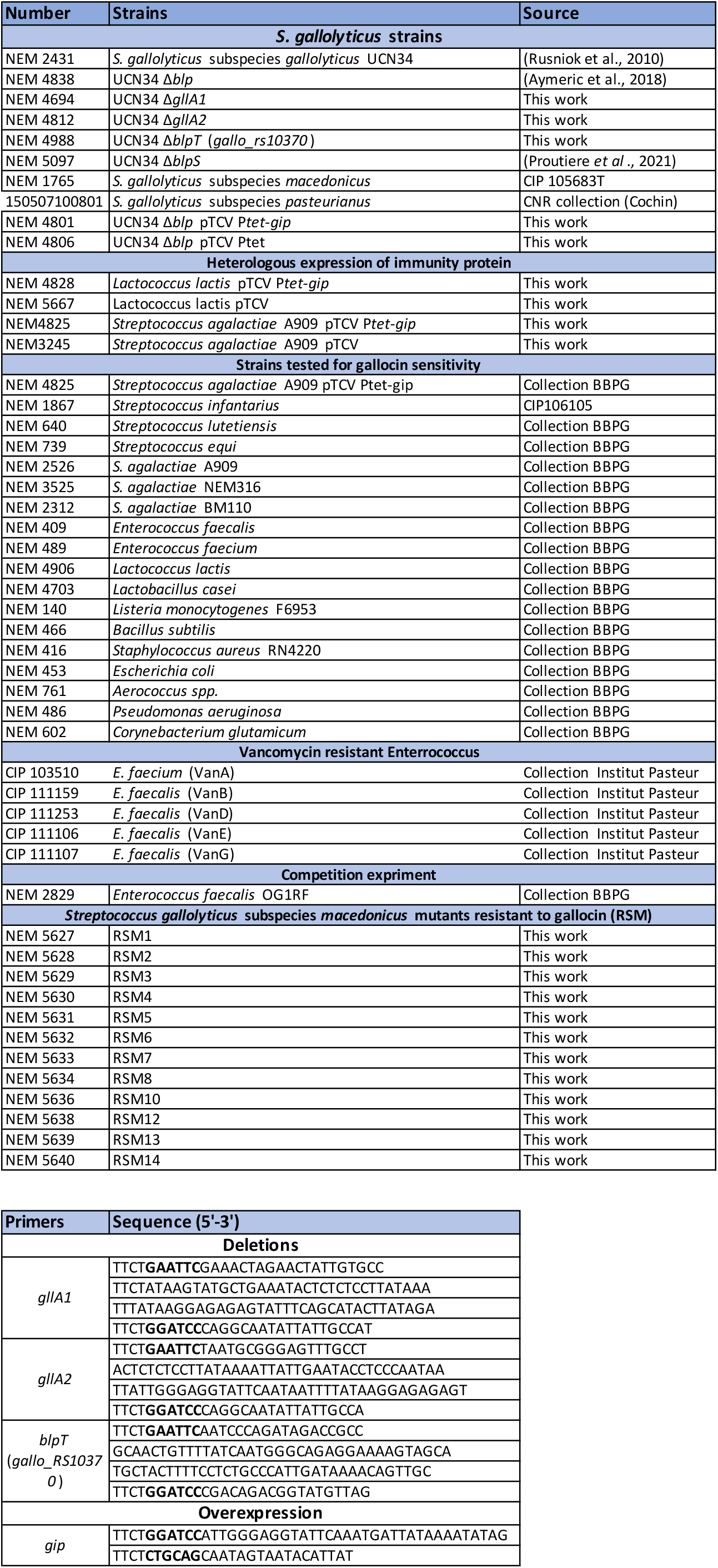
List of strains and primers.

The WalRK TCS is known as the master regulator of cell wall homeostasis, cell membrane integrity, and cell division processes in gram-positive bacteria (11). In streptococci, response regulator WalR (VicR) but not the histidine kinase WalK (VicK) is essential. Consistent with this, the 2 mutations observed in WalR were single amino acid substitutions (RSM6 Ala_95_ to Val; RSM12 Arg_117_ to Cys) while 4 out of the 5 mutations in WalK led to a frameshift or the appearance of a STOP codon (Fig. S6).

Interestingly, three other mutants (RSM7, RSM8 and RSM10) mapped in a single gene encoding a putative cell-wall binding protein with a C-terminal lysozyme-like domain. Two mutants (RSM7 and RSM8) exhibited frameshift mutations leading to the appearance of a premature STOP codon, and the last one (RSM10) a substitution of the putative key catalytic residue of the lysozyme-like domain (E_137_ to K, Fig. S6).

Thus, we hypothesized that peptidoglycan alterations in these mutant strains could explain the resistance to gallocin A. To test this hypothesis, we labelled peptidoglycan with the fluorescent lectin Wheat Germ Agglutinin (WGA-488) and imaged the mutants with conventional fluorescence microscopy. As shown in Fig. 7, most gallocin A resistant mutants, including all WalKR mutants, exhibit abnormal morphology and formed small aggregates as compared to the typical *SGM* WT linear chain of 2-5 cells. Cell morphology defects and peptidoglycan alterations were also detected in the 2 mutants which do not share common mutations with the other mutants (RSM 3 and 13, Fig. 7).

**Fig. 7.**
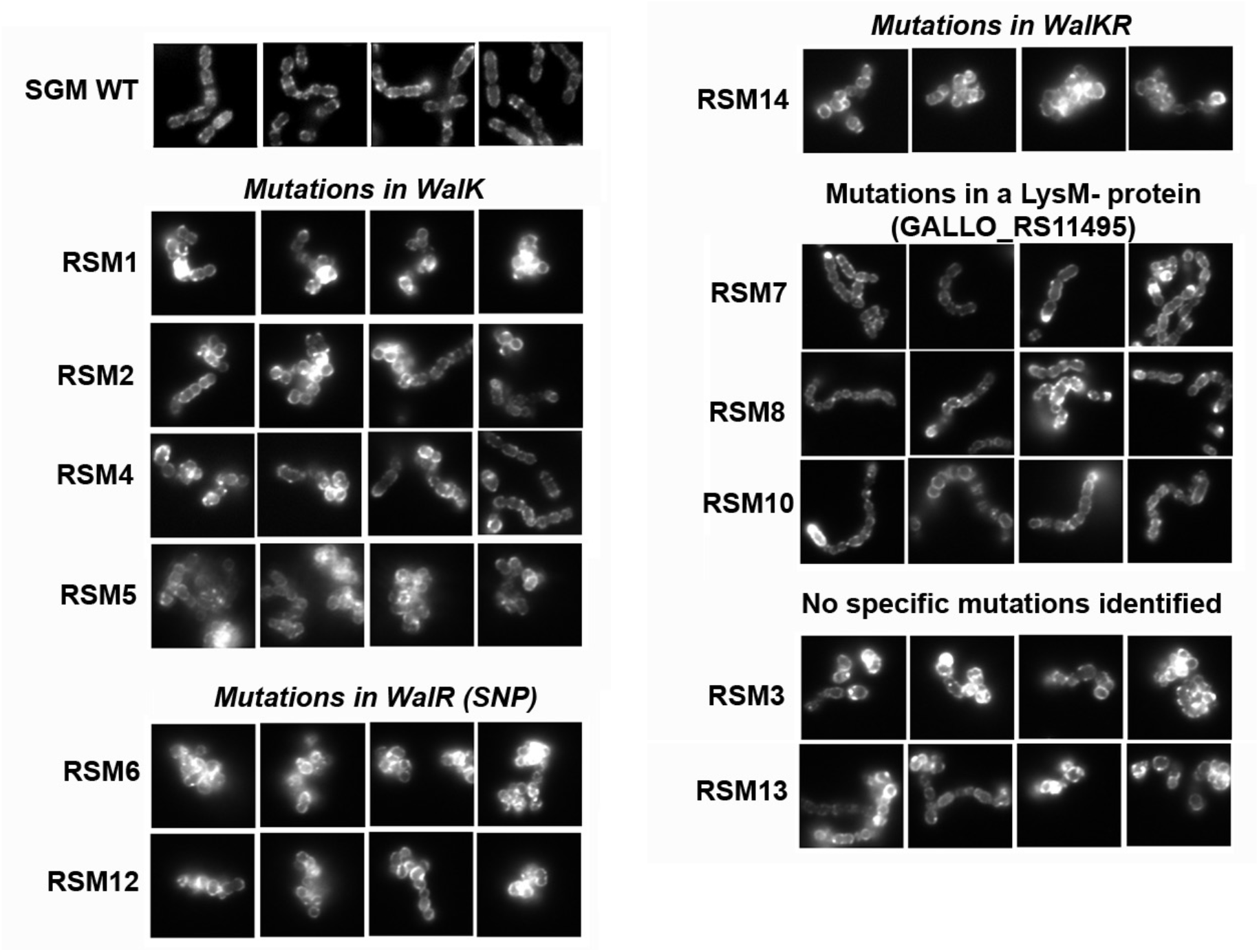
Gallocin A-resistant mutants (RSM) forms aggregates and exhibit morphological defects as compared to the parental gallocin A-sensitive strain *SGM*. Epifluorescence microscopy images of *SGM* WT and RSM 1 to 12 labelled with the Wheat Germ Agglutinin-488, a fluorescent peptidoglycan dye.

Taken together, these results suggest that alteration of the peptidoglycan structure could lead to gallocin A resistance, either by blocking its access to the membrane or by the formation of cell aggregates. It is worth noting that RSM mutants’ resistance to gallocin A was intermediate and that no potential membrane receptor for gallocin A peptides was identified.

## DISCUSSION

Gallocin A is a class IIb bacteriocin secreted by *Streptococcus gallolyticus subsp. gallolyticus* (*SGG*) to outcompete indigenous gut *Enterococcus faecalis* (*EF*) in tumoral conditions only (1). Mechanistically, gallocin A activity was found to be enhanced by higher concentrations of secondary bile acids found in tumoral conditions (1). Another proof-of-concept study showed that *EF* carrying the conjugative plasmid pPD1 expressing bacteriocin was able to replace indigenous enterococci lacking pPD1 (11). The rise of antimicrobial resistance combined with the recognized roles in health of gut microbiota homeostasis has attracted a renewed interest in the role of bacteriocins in gut colonization and their use as potential tools for editing and shaping the gut microbiome (12).

We show here that gallocin A, like many class IIb bacteriocins, only kills closely related species belonging to the Streptococcaceae and Enterococcaceae family. Interestingly, gallocin A can kill *Enterococcus faecium*, a commensal bacterium contributing largely to the transfer of antibiotic resistance in the microbiome and classified as high priority in the “WHO priority pathogens list for R&D of new antibiotics”. Taken together, these results highlight the potential of using bacteriocins such as gallocin A to fight antibiotic resistance and to cure bacterial infections with a lower impact on the gut microbiota due to their narrow spectrum of action.

Both GllA1 and GllA2 are synthesized as pre-peptides with an N-terminal leader sequence which is cleaved during export after a GG motif via a specific ABC transporter, BlpAB, to produce the extracellular mature active peptides (5). Experimental determination of the molecular mass of GllA1 and GllA2 by LC/MS fits with a cleavage after the GG motif present in the leader sequence and indicates the presence of an intramolecular disulfide bond in GllA1 and GllA2. Moreover, reduction of these disulfide bonds abrogates gallocin A antimicrobial activity. ColabFold modeling of GllA1 and GllA2 indicates that the N-terminal leader sequence is unstructured and that the mature GllA1 and GllA2 share a similar structural fold with two antiparallel α-helices forming a hairpin stabilized by an intramolecular disulfide bond. To our knowledge, this is the first report of an intramolecular disulfide bond in class IIb bacteriocin peptides. Most class IIb peptides, including the well described lactococcin G, the plantaricin EF, the plantaricin JK and the carnobacteriocin XY (CbnXY) (13–16), do not contain cysteine residues in their primary amino acid sequences. Consistently, the peptides constituting these 4 well-known bacteriocins are composed of only one main alpha-helix, and therefore do not require any disulfide bond to stabilize their tri-dimensional structures. Recently, gallocin D was identified in a very peculiar strain SGG LL009, isolated from raw goat milk in New Zealand (2). Gallocin D is a two-peptide bacteriocin homologous to infantaricin A secreted by *Streptococcus infantarius*, a member of the *Streptococcus bovis group* (2). Of note, the peptides of the 4 well described two-peptide bacteriocins discussed above and of gallocin D are much smaller in size (about 30 amino acids long) than the gallocin A peptides (2). In addition, gallocin A peptides are less positively charged (1 positively charged amino acid in GllA1, 2 in GllA2), while the highly positively charged C-terminus of lactococcin G α-peptide is thought to contribute to the anchoring of the peptide to the membrane, thanks to the transmembrane potential (negative inside) (13, 17).

A few other class IIb bacteriocins, such as brochocin C, thermophilin 13 and ABP-118 (18–21), were found to share similar structural properties with gallocin A peptides (longer peptides, few positively charged amino acids and two cysteine residues in each peptide located close to N-/C-terminus). Alphafold modelling of these peptides showed that their putative structures resemble those of GllA1 and GllA2, with two-antiparallel alpha-helices. Disulfide bonds between the cysteines of the 2 helices were also predicted in 5 out of the 6 peptides (Fig. S7). BrcB, the peptide without predicted disulfide bond, was also the one with the worse IDDT score, suggesting that the prediction might not be accurate. In conclusion, gallocin A, as well as other class IIb bacteriocins such as brochocin C, thermophilin 13 and ABP-118, might represent a subgroup in class IIb bacteriocins which differs in structure, and potentially in their mode of action from the other well-known class IIb bacteriocins.

Finally, gallocin A resistance was studied through whole-genome sequencing of 12 spontaneous resistant mutants derived from the highly sensitive strain *S. gallolyticus* subsp. *macedonicus* CIP105683T. Previously, this method allowed the identification of UppP as a membrane receptor required for lactococcin G activity (7). Unlike this previous study, we did not find a common gene mutated in our 12 resistant mutants (RSM), suggesting that gallocin A does not require the presence of a specific receptor. This is in agreement with our data showing that gallocin A can permeabilize lipid vesicles composed of two phospholipids (phosphatidylcholine and phosphatidylglycerol). The majority of RSM mutants exhibited mutations in the genes encoding a regulatory two-component system sharing strong homologies with WalKR (also known as VicKR and YycGF). This two-component system, originally identified in *Bacillus subtilis*, is very highly conserved and specific to low GC% Gram-positive bacteria, including several pathogens such as *Staphylococcus aureus* (22, 23). Several studies have unveiled a conserved function for this system in different bacteria, including several streptococcal pathogens, defining this signal transduction pathway as a master regulatory system for cell wall metabolism (23). Consistent with the potential defect in cell-wall synthesis, these mutants showed morphological abnormalities and cell-division defects. Similar observations have been reported in *Staphylococcus aureus* (24–26) where mutations in *walK* were shown to confer intermediate resistance to vancomycin and daptomycin.

Three mutants displayed independent mutations in a small protein (197 amino acids) of unknown function containing an N-terminus LysM-peptidoglycan binding domain and a C-terminus lysozyme-like domain. The lysozyme-like domain, which is about fifty amino acids long, was originally identified in enzymes that degrade the bacterial cell-walls. Interestingly, the mutations in RSM7, RSM8, RSM10 mutant all mapped within the lysozyme-like domain, suggesting a potential alteration of the cell-wall in these mutants. Finally, the last two last mutants (RSM3 and RSM13) carrying mutations in other genes than in *walRK* exhibited the same morphology defects associated with gallocin A resistance.

To conclude, it is worth highlighting that the 12 mutants were only partially resistant to gallocin A. Most RSM mutants form bacterial aggregates which probably contributes to their resistance to gallocin A, just as biofilms are more resistant to antibiotics. No specific membrane receptor could be identified for gallocin A. Interestingly, it has also been suggested that thermophilin 13, another class IIb bacteriocin that shares putative structural similarity with gallocin A (18), does not require any specific receptor for its activity. However, the different level of susceptibility to gallocin A within a given species, as demonstrated for three Group B *Streptococcus* strains (A909 > BM110> NEM316), as well as its narrow-spectrum mode of action indicate that unidentified bacterial factors can modulate gallocin A sensitivity. It will also be important in the future to identify the direct bacterial targets of gallocin A in the murine colon using global 16S DNA sequencing in normal and tumoral conditions.

## MATERIALS AND METHODS

### Cultures, bacterial strains, plasmids and oligonucleotides

*Streptococci* and *Enterococci* used in this study were grown at 37°C in Todd-Hewitt broth supplemented with 0.5% yeast extract (THY) in standing filled flasks. When appropriate, 10 μg/mL of erythromycin were added for plasmid maintenance.

Plasmid construction was performed by: PCR amplification of the fragment to insert in the plasmid with Q5^®^ High-Fidelity DNA Polymerase (New England Biolabs), digestion with the appropriate FastDigest restriction enzymes (ThermoFisher), ligation with T4 DNA ligase (New England Biolabs) and transformation in commercially available TOP10 competent *E. coli* (ThermoFisher). *E. coli* transformants were cultured in Miller’s LB supplemented with 150 μg/mL erythromycin (for pG1-derived plasmids) or 50 μg/mL kanamycin (pTCV-derived plasmid). Verified plasmids were electroporated in *S. agalactiae* NEM316 and mobilized from NEM316 to *SGG* UCN34 by conjugation as described previously (27). pTCV-derived plasmids were electroporated in *Lactococcus lactis* NZ9000. Strains, plasmids and primers used in this study are listed in Table 1. The wide range of bacteria tested *in vitro* for their resistance or sensitivity to gallocin A antimicrobial activity come from our laboratory repository and were cultured in their optimal media and conditions.

### Construction of markerless deletion mutants in SGG UCN34

In-frame deletion mutants were constructed as described previously (27). Briefly, the 5’ and 3’ region flanking the region to delete were amplified and assembled by splicing by overlap extension PCR and cloned into the thermosensitive shuttle vector pG1. Once transformed in UCN34, the cells were cultured at 38°C with erythromycin to select for the chromosomal integration of the plasmid by homologous recombination. About 4 single cross-over integrants were serially passaged at 30 °C without antibiotic to facilitate the second event of homologous recombination and excision of the plasmid resulting either in gene deletion or back to the WT (bWT). In-frame deletions were identified by PCR and confirmed by DNA sequencing of the chromosomal DNA flanking the deletion.

### Gallocin A production assays

Briefly, one colony of the indicator strain, here *Streptococcus gallolyticus subps. macedonicus* (*SGM*), was resuspended in 2 mL THY, grown until exponential phase, poured on a THY agar plate, the excess liquid was removed and left to dry under the hood for about 20 min. Using sterile tips, 5-mm-diameter wells were dug into the agar. Each well was then filled with 80 μL of filtered supernatant from 5 h cultures (stationary phase) of *SGG* UCN34 WT or otherwise isogenic mutant strains and supplemented with Tween 20 at 0.1% final concentration. Inhibition rings around the wells were observed the following morning after overnight incubation at 37°C.

### Competition experiments

*SGG* strains were inoculated from fresh agar plate at initial DO_600_ of 0.1 together with *E. faecalis* OG1RF in THY medium and incubated for 4 h at 37°C in micro aerobiosis. After 4 h of co-culture, the mixed cultures were serially diluted and plated on Enterococcus agar-selective plates (BD Difco). On these plates, *SGG* exhibits a pale pink color while *E. faecalis* exhibits a strong purple color. CFU were counted the next morning to determine the final concentration in CFU/mL in each test sample.

### Analysis of gallocin A peptides by LC-MS

*Sgg* UCN34 was grown in 500 mL of sterile THY supplemented with 5 nM synthetic GSP at 37 °C with 5% CO_2_ for 12-16 h. The cultures were centrifuged at 4,000 x g for 20 min and the supernatant was filtered through a sterile 0.22 μm polyethersulfone (PES) filter. Ammonium sulfate was added to the filtered supernatants to give a 20% (wt/vol) concentration and mixed by inversion until all ammonium sulfate salts went into solution. The solution was stored at 4 °C for 1 h, followed by centrifugation at 4,000 × g for 20 min. The supernatants were discarded, and the remaining pellet was dissolved in 100 mL DI water and placed in a 3 kDa MWCO dialysis tube. The dialysis tube was placed in a 500 mL graduated cylinder containing distilled water and a stir bar. Dialysis was performed for 4 h with changing of DI water every hour. The material in the dialysis tube was then lyophilized. A 5 mg/mL solution of the lyophilized material was prepared in 75:25 (H_2_O:ACN) and 50 μL were injected into an Agilent Technologies 6230 time of flight mass spectrometer (an HRMS system) with the following settings for positive electrospray ionization (ESI+) mode: capillary voltage = 3,500 V; fragmentor voltage = 175 V; skimmer voltage = 65 V; Oct 1 RF Vpp = 750 V; gas temperature = 325 °C; drying gas flow rate = 0.7 L/min; nebulizer; 25 lb/in2; acquisition time = 17.5 min. An XBridge C18 column (5 μm, 4.6 × 150 mm) was used for the LC-MS analysis.

### Membrane permeabilization assays

These assays were performed as described previously (28). Briefly, ANTS (fluorophore probe) and DPX (quencher) were encapsulated into large unilamellar vesicles (LUVs) to monitor membrane permeabilization induced by peptides. The LUVs were prepared at a concentration of 10 mM lipid at a POPC:POPG molar ratio of 8:2 containing 20 mM ANTS and 60 mM DPX. The multilamellar vesicle suspension was extruded through 0.4- and 0.2-μm polycarbonate filters to produce LUVs of 200 ± 30 nm in diameter, as measured by DLS. The unencapsulated ANTS and DPX were removed by gel filtration through a Sephadex G-25 column 5 mL (Cytiva, USA). For permeabilization assays, LUVs were incubated in buffer at 0.45 mM lipids at 25 ⍰C in a 101-QS cuvette (Hellma, France) and under constant stirring. The excitation wavelength was set to 390 nm and the emission of ANTS was continuously measured at 515 nm. The maximum intensity of permeabilization, corresponding to the maximum recovery of ANTS fluorescence was measured after addition of 0.12% (2 mM) of Triton X100.

### Generation of gallocin resistant mutants

In order to generate gallocin resistant mutants, we concentrated *SGG* supernatant 200 times by precipitation with 20% of ammonium sulfate. By serial 2-fold dilutions, we showed that this supernatant was approximatively 64 times more concentrated than the original supernatant (Fig. S5A). Fourteen resistant mutants (named RSM1 to 14) of *S. gallolyticus subsp. macedonicus* parental strain CIP105683T, the species showing the highest sensitivity to gallocin A, were selected on THY agar plates containing 10% of this concentrated supernatant. Twelve of them were confirmed to be gallocin resistant by growth in THY supplemented with the supernatant of SGG WT, containing gallocin, and 0.01% of Tween20, which is necessary for gallocin A activity (Fig. S4B and C). As an important control, the same experiment was performed after precipitation of the Δ*blp* supernatant that does not produce gallocin A. *SGM* WT was re-isolated on this plate and a single colony was stocked and sequenced with the RSM mutants as described below.

### Sequencing and SNP localization

Whole-genome sequencing of the control *SGM* WT, re-isolated from Δ*blp* plate as described in the section above, and of RSM mutants was performed using Illumina technology and compared with the genome of the parental strain *SGM* CIP105683T that was *de novo* assembled using PacBio sequencing. The assembly was performed with Canu 1.6. (29) leading to a main chromosome of 2,210,410bp and a plasmid of 12729bp (HE613570.1). The annotation was subsequently made with Prokka (30) before a variant calling was performed using the Sequana (31) variant calling pipeline. Of note, variants were called with a minimum frequency of 10% and a minimum strand balance of 0.2. Many mutations, probably due to the different method used for the sequencing of the reference sequence, were present in the control *SGM* WT strain used as control and the RSM mutants. Therefore, only RSM specific mutations occurring at a frequency>0.5 as compared to control SGM WT were taken into account for this analysis and are shown in Table 2.

**Table 2:**
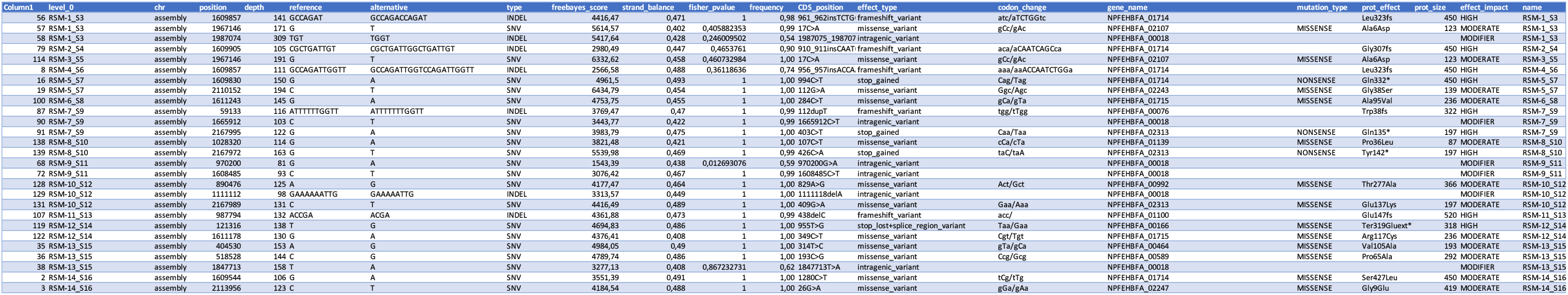
Single nucleotide polymorphisms detected in RSM mutants as compared to the control *SGM* WT.

## ACKNOWLEDGEMENTS

We would like to thank particularly Tarek Msadek for careful and critical reading of the manuscript. This study has been funded by the Institut National contre le Cancer (INCA) PLBIO 16-025 attributed to S. Dramsi and from the French Government’s Investissement d’Avenir program, Laboratoire d’Excellence “Integrative Biology of Emerging Infectious Diseases (grant n° ANR-10-LABX-62-IBEID). This study was also funded by the National Science Foundation (NSF, CHE-1808370, to Y.T).

## SUPPLEMENTARY FIGURE LEGENDS

**Fig. S1. Gallocin A is active in a broad range of pH and heat-stable**

**A)** Agar diffusion assay to test gallocin activity from supernatants of UCN34 WT and Δ*blp* at different pH against *SGM*, a gallocin A-sensitive bacterium. The initial supernatant from an overnight culture (left well) had a pH of 5.4. pH was then adjusted to 2-12 using HCl or NaOH. **B)** Supernatant was heated at 80°C for indicated times and tested as in (A).

**Figure S2: Gallocin A spectrum of action**

Agar diffusion assay using UCN34 WT and Δ*blp* supernatant against various bacterial species.

**Fig. S3: LC-MS analysis of the two components comprising gallocin A indicating that both peptides contain a disulfide bridge**.

Left Panel: Top – GllA2 structure, molecular formula and molecular weight (with disulfide bridge); Middle – Mass Spectrum showing the MH_3_^+3^/3 and MH_4_^+4^/4 masses observed for GllA2; Bottom – LC chromatogram showing the peak where GllA2 was detected. Right Panel: Top – GllA1 structure, molecular formula and molecular weight (with disulfide bridge); Middle – Mass Spectrum showing the MH_2_^+2^/2 and MH_3_^+3^/3 masses observed for GllA1; Bottom – LC chromatogram showing the peak where GllA1 was detected.

**Fig. S4: Structural models of GllA1, GllA2 and GIP alone or in complex with each over**.

All representations are colored with predicted lDDT from a score of 30% (red) to 100% (blue). The disulfide bond is visible in stick representation for GllA1 and GllA2.

**Fig. S5: Obtention of 12 spontaneous mutants (RSM) resistant to gallocin A as compared to the parental sensitive strain *SGM***.

**A)** Agar diffusion assay against *S. macedonicus* using serial two-fold dilutions of *Sgg* supernatant concentrated (SN 200X) or not (SN 1X) by ammonium sulfate precipitation. **B)** and **C)** Growth curves for the 12 gallocin A-resistant mutants in the presence or absence of gallocin A (THY medium supplemented with 30% of SGG WT/Δ*blp* supernatant)

**Fig. S6: Amino acid changes in WalK, WalR or Aggregation Promoting Factor (APF) in RSM mutants as compared to the parental SGM (WT)**

Comparison of the primary amino acid sequence of WalK (A), WalR (B), and the aggregation promoting factor (C) found in RSM mutants to their WT counterpart. Putative domains of these proteins, identified by BLAST, are shown in red (HATPase_C: <u>smart00387</u>; REC: <u>cd17614</u>; Helix-turn-helix: <u>pfam00486</u>; LysM: <u>cd00118</u> and Lysozyme-like: <u>cd13925</u>). Putative residues important for the protein activity, identified by BLAST, are indicated by a dark arrowhead.

**Fig. S7: Putative structure of two-component bacteriocins**

Structural models of mature forms of the two peptides composing the two-component bacteriocins ABP118, Brochocin C and Thermophilin 13 using ColabFold. The amino acid sequence following the first glycine doublet (in bold) was considered as the mature form of the peptides. Uniprot accession numbers: ABP118: Q8KWI0; Q8KWH9. Brochocin C: O85756; O85757. Thermophilin 13: O54454; O54455. All representations are colored with predicted lDDT from a score of 30% (red) to 100% (blue). The disulfide bond is visible in stick representation for all the peptides except BrcB.

## Notes

### Competing Interest Statement

The authors have declared no competing interest.

### Summary of Updates

Title and abstract have been revised !

