## Supplemental Figures 1 to 7 for "Gallocin A, an atypical two-peptide bacteriocin with intramolecular disulfide bonds required for activity"

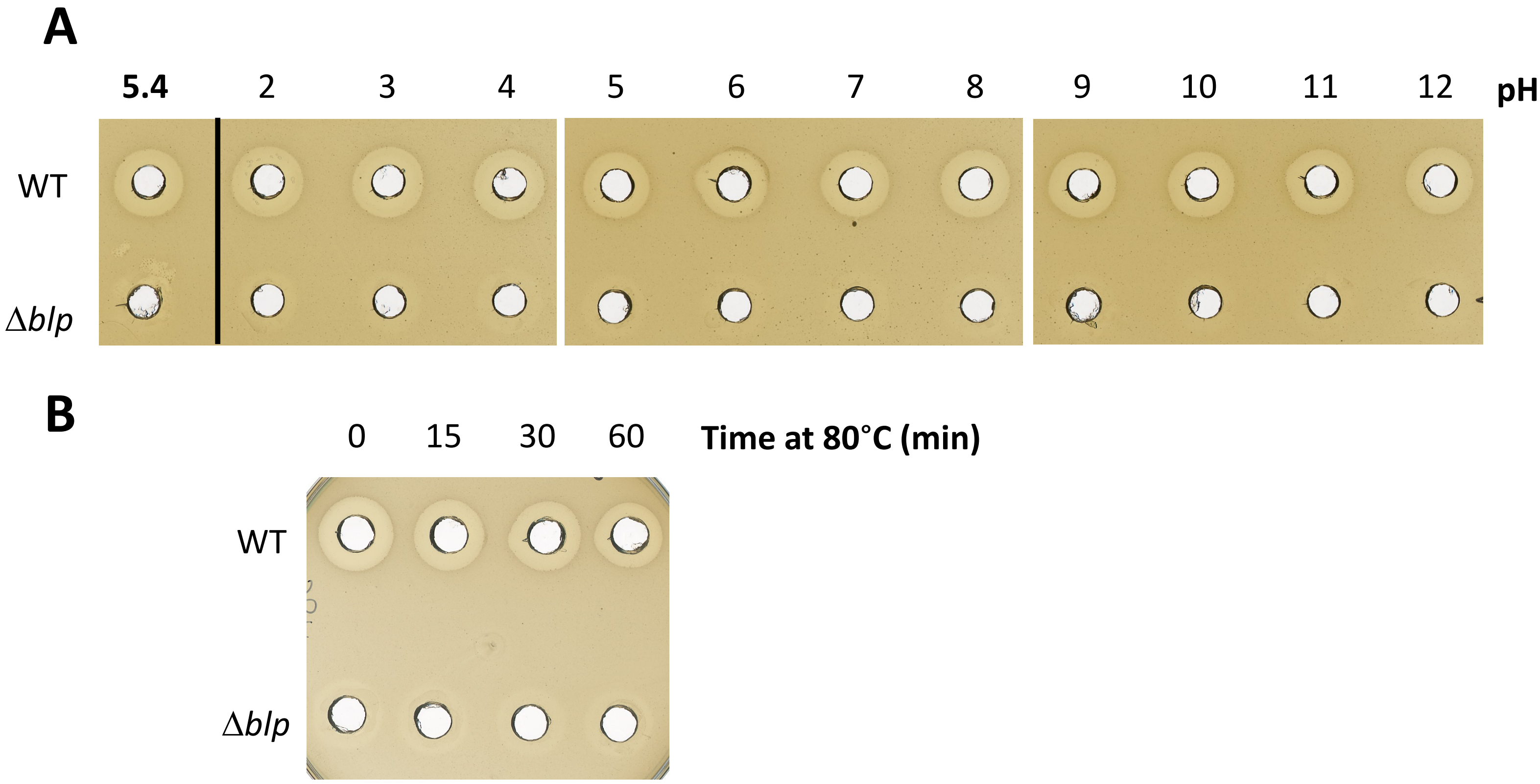


**Figure S1 : Gallocin A is active in a broad range of pH and heat-stable**

A) Agar diffusion assay to test gallocin activity from supernatants of *SGG* WT and Δ*blp* at different pH against gallocin A- sensitive *SGM.* The initial supernatant from an overnight culture (left well) had a pH of 5.4. pH was then adjusted to 2-12 using HCl or NaOH. B) Supernatant was heated at 80°C for indicated times and tested as in (A).

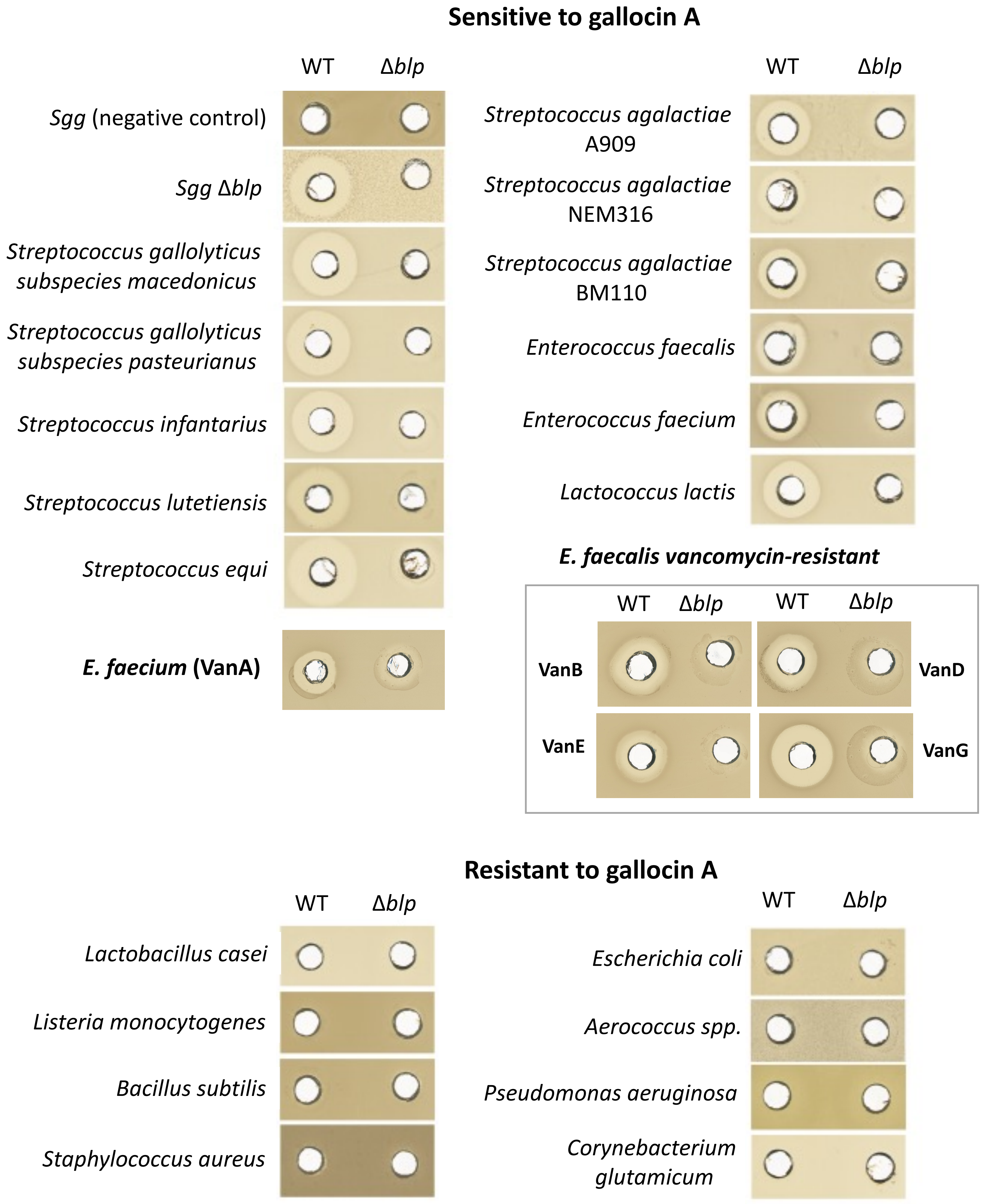


**Figure S2: Gallocin A spectrum of action**

Agar diffusion assay using *Sgg* WT and Δ*blp* supernatant against various bacterial species

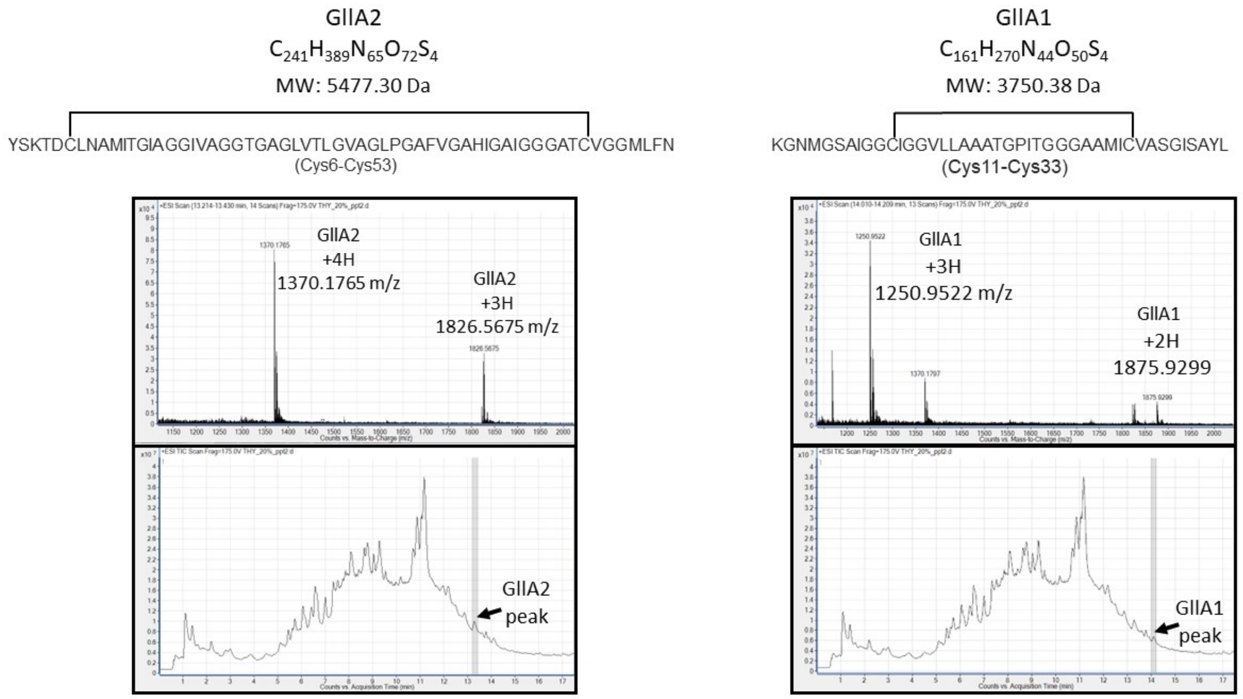


**Figure S3.** LC-MS analysis of the two components comprising gallocin exhibiting that both peptides contain a disulfide bridge. Left Panel: Top – GllA2 structure, molecular formula and molecular weight (with disulfide bridge); Middle – Mass Spectrum showing the $MH_3^{+3}/3$ and $MH_4^{+4}/4$ masses observed for GllA2; Bottom – LC chromatogram showing the peak where GllA2 was detected. Right Panel: Top – GllA1 structure, molecular formula and molecular weight (with disulfide bridge); Middle – Mass Spectrum showing the $MH_2^{+2}/2$ and $MH_3^{+3}/3$ masses observed for GllA1; Bottom – LC chromatogram showing the peak where GllA1 was detected.

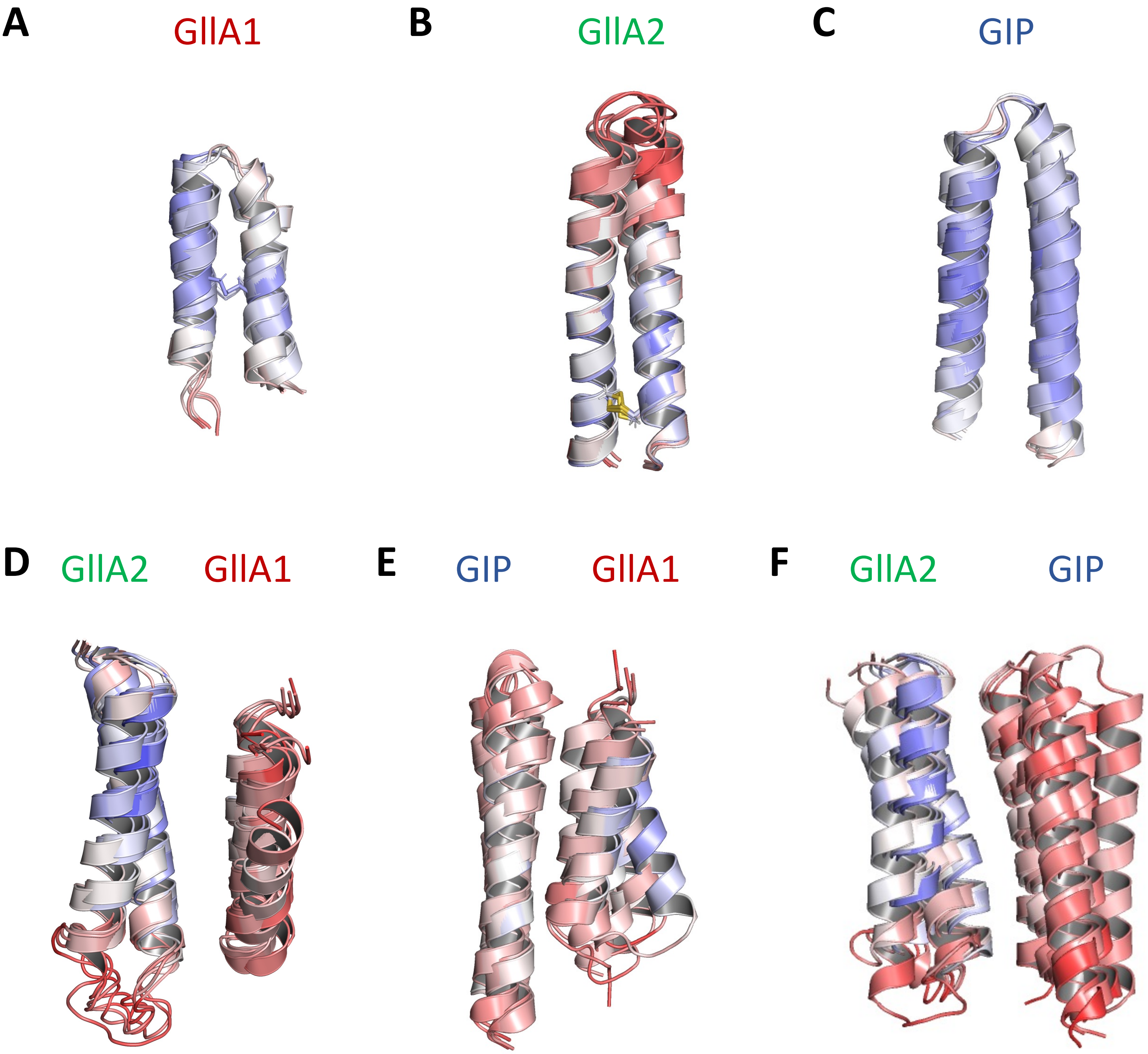


**Figure S4: Putative structures of GIIA1, GIIA2 and GIP alone or in complex with each over.** All representation are colored with predicted lDDT from a score of 30% (red) to 100% (blue). The disulfide bond is visible in stick representation for GIIA1 and GIIA2.

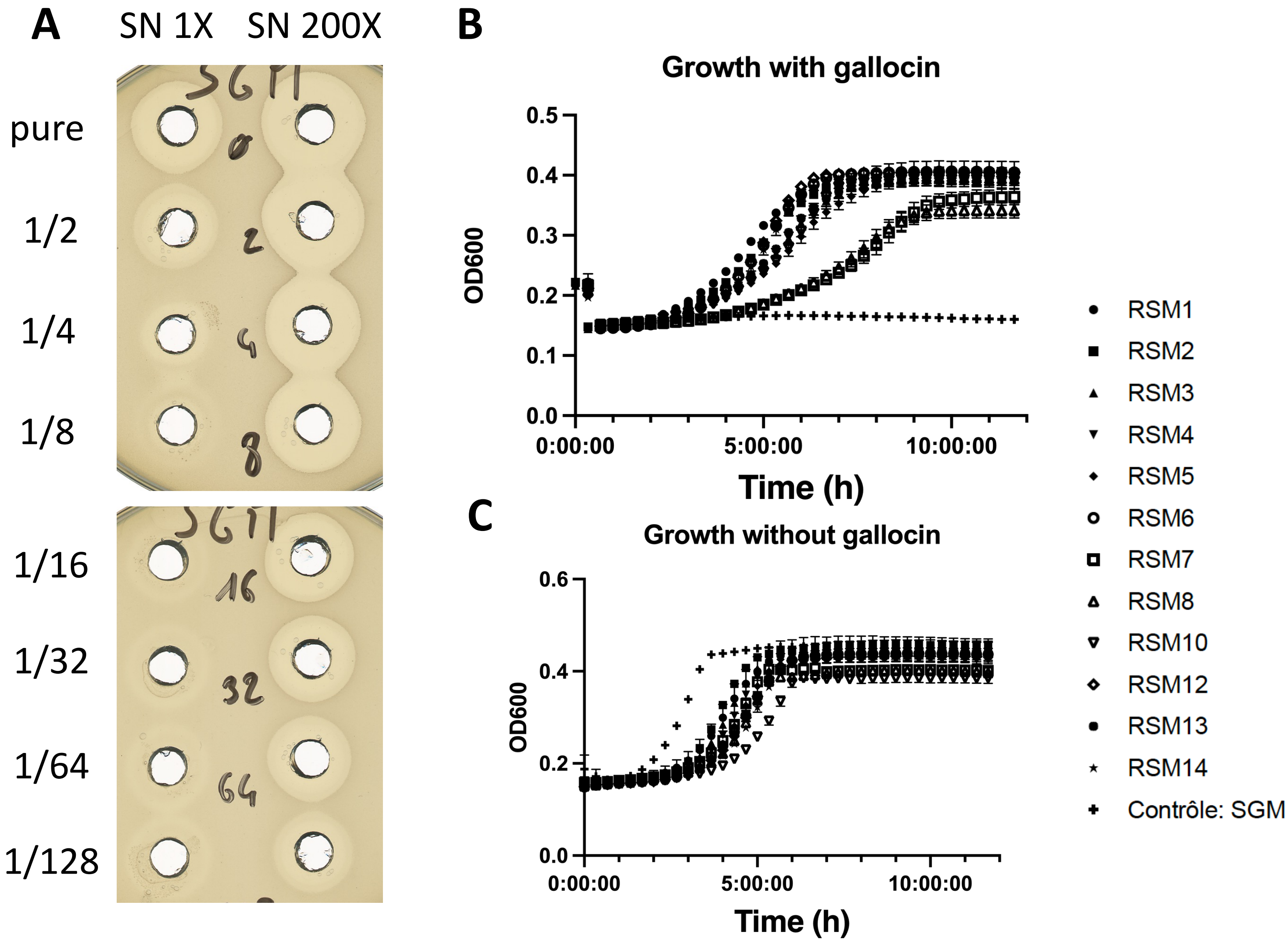


**Figure S5: Generation of *SGM* mutant resistant to gallocin A named RSM1 to 14**

A) Agar diffusion assay against *S. macedonicus* using serial two-fold dilutions of *Sgg* supernatant concentrated (SN 200X) or not (SN 1X) by ammonium sulfate precipitation. B) and C) Growth curve of RSM 1 to 12 in presence or absence of gallocin (THY medium supplemented with 30% of Sgg WT/Δ*blp* supernatant)

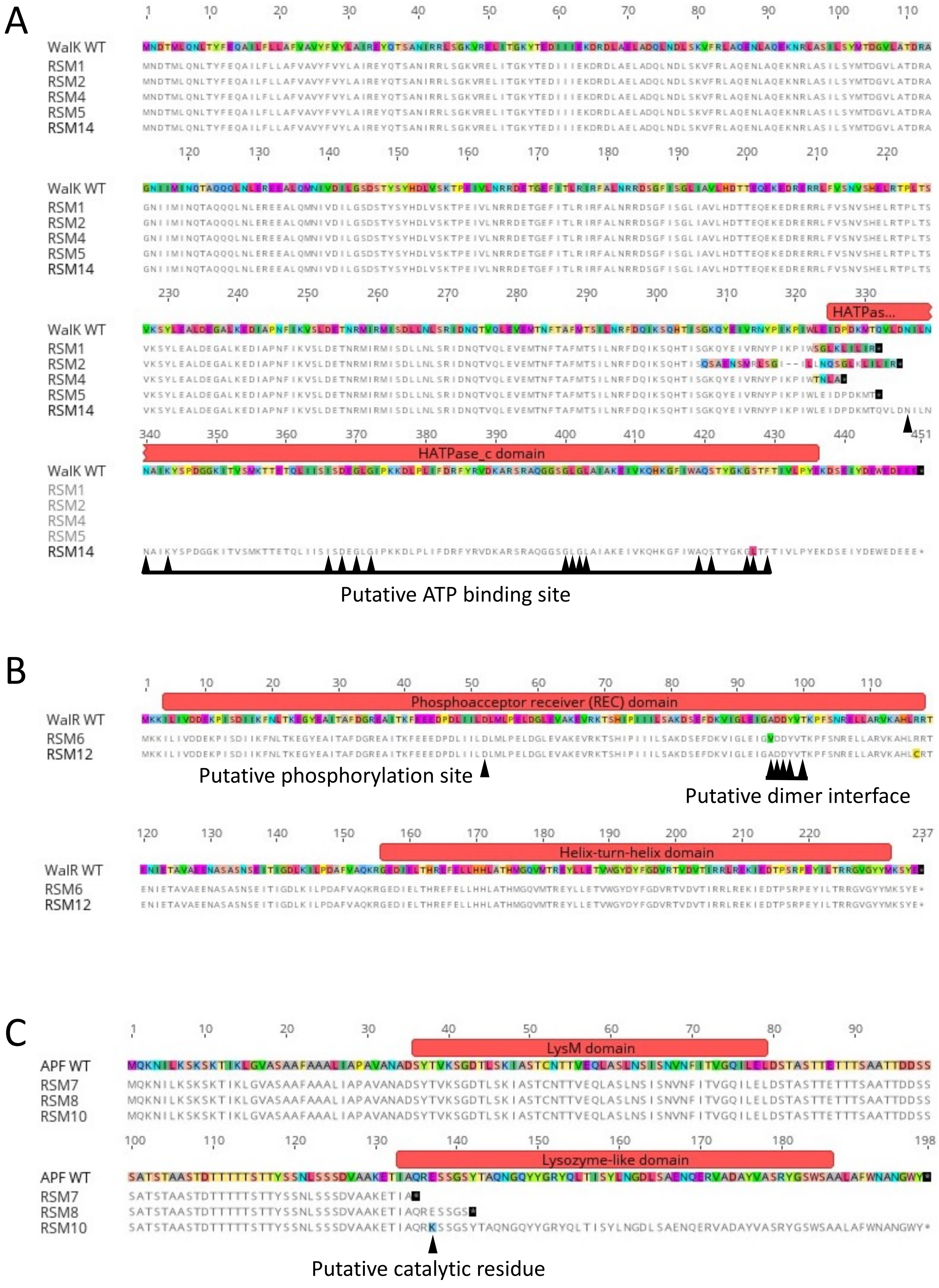


**Figure S6: Amino acid changes in WalK, WalR or Aggregation Promoting Factor (APF) in RSM mutants as compared to the parental SGM (WT)**

Comparison of the amino acid sequences of WalK (A), WalR (B), and the aggregation promotig factor found in RSM mutants to their WT counterpart. Putative domains of these proteins, identified by BLAST, are shown in red (HATPase_C: smart00387 ; REC: cd17614 ; Helix-turn-helix: pfam00486 ; LysM: cd00118 and Lysozyme-like: cd13925). Putative residues important for the protein activity, identified by BLAST, are indicated by dark arrowheads.

**A**

ABP118-α

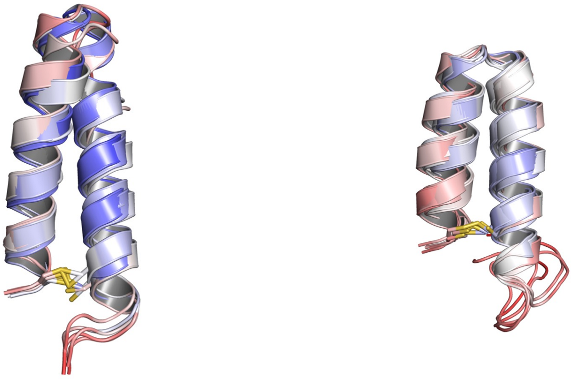

ABP118-β

**ABP118-α**: MMKEFTVLTECELAKVDGG**KRGPNCVGNFLGGLFAGAAAGVPLGPAGIVGGANLGMVGGALTCL**
**ABP118-β**: MKNLDKRFTIMTEDNLASVNGG**KNGYGGSGNRWVHCGAGIVGGALIGAIGGPWSAVAGGISGGFTSCR**

**B**

BrcA

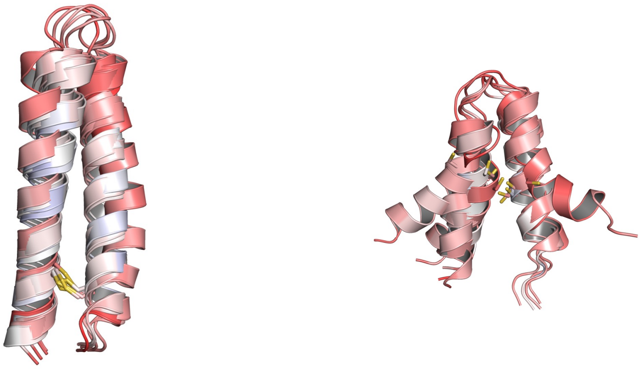

BrcB

**BrcA**: MHKVKKLNNQELQQIVGG**YSSKDCLKDIGKGIGAGTVAGAAGGGLAAGLGAIPGAFVGAHFGVIGGSAACIGGLLGN**
**BrcB**: MKKELLNKNEMSRIIGG**KINWGNVGGSCVGGAVIGGALGGLGGAGGGCITGAIGSIWDQW**

**C**

ThmA

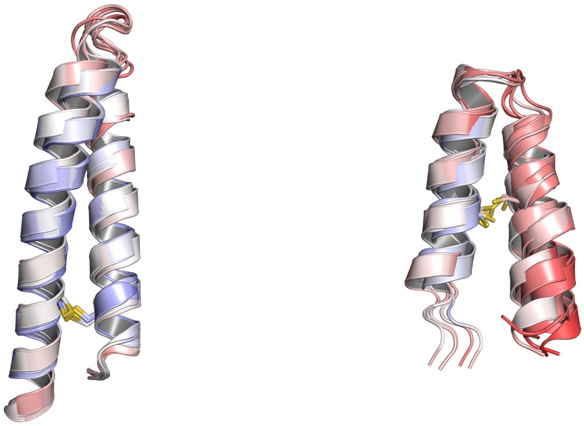

ThmB

**ThmA**:
MNTITICKFDVLDAELLSTVEGG**YSGKDCLKDMGGYALAGAGSGALWGAPAGGVGALPGAFVGAHVGAIAGGFACM GGMIGNKFN**
**ThmB**: MKQYNGFEVLHELDLANVTGG**QINWGSVVGHCIGGAIIGGAFSGGAAAGVGCLVGSGKAIINGL**

**Figure S7: Putative structure of two-component bacteriocins**

Predicted structure of mature forms of the two peptides composing the two-component bacteriocins ABP118, Brochocin C and Thermophilin 13 using Colabfold. The amino acid sequence following the first glycine doublet (in bold) was considered as the mature form of the peptides. Uniprot accession numbers: ABP118: Q8KWI0 ; Q8KWH9. Brochocin C: O85756 ; O85757. Thermophilin 13: O54454 ; O54455. All representation are colored with predicted lDDT from a score of 30% (red) to 100% (blue). The disulfide bond is visible in stick representation for all the peptides except BrcB.
